# Postsynaptic synucleins mediate vesicular exocytosis of endocannabinoids

**DOI:** 10.1101/2021.10.04.462870

**Authors:** Eddy Albarran, Yue Sun, Yu Liu, Karthik Raju, Ao Dong, Yulong Li, Sui Wang, Thomas C. Südhof, Jun B. Ding

**Affiliations:** Neurosciences Graduate Program; Department of Neurosurgery; Department of Molecular and Cellular Physiology, and Howard Hughes Medical Institute; State Key Laboratory of Membrane Biology, Peking University School of Life Sciences, Beijing, China; PKU-IDG/McGovern Institute for Brain Research, Beijing, China; Peking-Tsinghua Center for Life Sciences, Academy for Advanced Interdisciplinary Studies, Peking University, Beijing, China; Chinese Institute for Brain Research, Beijing, China; Department of Ophthalmology; Department of Neurology and Neurological Sciences, Stanford University, Stanford, CA 94305, USA

## Abstract

Two seemingly unrelated questions have long motivated studies in neuroscience: How are endocannabinoids, among the most powerful modulators of synaptic transmission, released from neurons? What are the physiological functions of synucleins, key contributors to Parkinson’s Disease? Here, we report an unexpected convergence of these two questions: Endocannabinoids are released via vesicular exocytosis from postsynaptic neurons by a synuclein-dependent mechanism. Specifically, we find that deletion of all synucleins selectively blocks all endocannabinoid-dependent synaptic plasticity; this block is reversed by postsynaptic expression of wildtype but not of mutant α-synuclein. Loading postsynaptic neurons with endocannabinoids via patch-pipette dialysis suppressed presynaptic neurotransmitter release in wildtype but not in synuclein-deficient neurons, suggesting that the synuclein deletion blocks endocannabinoid release. Direct optical monitoring of endocannabinoid release confirmed the requirement of synucleins. Given the role of synucleins in vesicular exocytosis, the requirement for synucleins in endocannabinoid release indicates that endocannabinoids are secreted via exocytosis. Consistent with this hypothesis, postsynaptic expression of tetanus-toxin light chain, which cleaves synaptobrevin SNAREs, also blocked endocannabinoid-dependent plasticity and release. The unexpected finding that endocannabinoids are released via synuclein-dependent exocytosis assigns a function to synucleins and resolves a longstanding puzzle of how neurons release endocannabinoids to induce synaptic plasticity.

---

α-Synuclein (α-Syn) is a small protein that, together with the closely related β- (β-Syn) and γ-Synucleins (γ-Syn), constitutes one of the most abundant proteins in the brain^1,2,3,4^. α-Syn plays a central role in Parkinson’s Disease (PD) pathogenesis since α-Syn mutations and multiplications cause PD^5,6,7,8,9^, genome-wide association studies link α-Syn to sporadic forms of PD^10,11^, and the brains of PD patients invariably contain Lewy bodies composed of α-Syn aggregates^12^. However, the physiological function of α-Syn, and that of other synucleins, remains largely unknown.

Synucleins possess a conserved N-terminal domain that binds to phospholipids^13,14,15^, underlying α-Syn’s affinity for membranes such as synaptic vesicles^16,17^. Overexpression of α-Syn *in vitro* and *in vivo* inhibits exocytosis, possibly through impairments in synaptic vesicle endocytosis, recycling, and dilation of the exocytotic fusion pore^17,18,19,20^. By contrast, deletion of α-Syn produces little to no effect on synaptic transmission, with α-Syn-KO mice exhibiting only slight reductions in dopamine (DA) levels and displaying modest behavioral phenotypes^21,22^. Moreover, synuclein double and triple knockout mice displayed no detectable changes in synaptic strength or short-term plasticity^23,24^. Thus, it has been difficult to reconcile α-Syn’s abundance and highly penetrant role in PD with its seemingly subtle endogenous function. Strikingly, even modest transgenic α-Syn overexpression completely prevents the lethality and neurodegeneration of CSPα KO mice^25^, suggesting an essential role for α-Syn in protection against neurodegeneration, which is counterintuitive given its causal involvement in PD.

The striatum, the input nucleus of the basal ganglia, is one of the most severely affected areas in PD, as the loss of DA signaling in the striatum and the degeneration of synapses on striatal spiny projection neurons (SPNs) greatly alter the striatal circuitry and underlie many of the motor and cognitive impairments observed in PD^26,27,28^. One particularly detrimental consequence of PD is the loss of endocannabinoid- (eCB-) dependent plasticity at corticostriatal synapses^29,30,31,32^, which is central to striatum-dependent learning and habit formation^33,34^. In eCB-dependent plasticity, eCBs are synthesized and released postsynaptically in an activity- and Ca^2+^-dependent manner. eCBs then retrogradely bind to presynaptic CB1 receptors (CB1Rs) to decrease the presynaptic release probability^35,36,37,38^. However, little is known about how eCBs are released from postsynaptic neurons. eCBs are amphiphilic molecules derived from phospholipids that are unlikely to diffuse passively out of the postsynaptic neurons and across the synaptic cleft^39,40^. Thus, how eCBs reach presynaptic CB1Rs during synaptic plasticity, an essential step to understanding striatal function, is unclear.

## Normal basal synaptic transmission in Syn-tKO mice

Given the strong association of corticostriatal dysfunction with PD, we directly measured basal corticostriatal synaptic transmission and eCB-dependent plasticity in α/β/γ-synuclein triple KO (Syn-tKO) mice. Previous reports suggested that α-Syn decreases neurotransmitter release by acting at presynaptic sites, with some studies showing increased synaptic transmission in single α-Syn KO mice^21,41^, whereas no such changes were detected in double^23^ or triple synuclein KO mice^24^. We therefore investigated if corticostriatal synaptic transmission was abnormal in Syn-tKO mice. Whole-cell patch clamp recordings from SPNs in acute slices of the dorsolateral striatum prepared from wildtype (WT) and Syn-tKO mice, combined with electrical stimulation of corticostriatal axons, allowed us to measure corticostriatal synaptic responses (Fig. 1a). We found no significant difference in the stimulus-response relationship between WT and Syn-tKO corticostriatal synapses (Fig. 1b). Because previous reports have shown that survival and behavioral deficits are revealed at older ages in Syn-tKO mice^24,42^, we also tested aged mice (16-18 months old). Again we observed no significant difference in synaptic strength between WT and Syn-tKO mice (Extended Data Fig. 1a,b).

**Figure 1.**
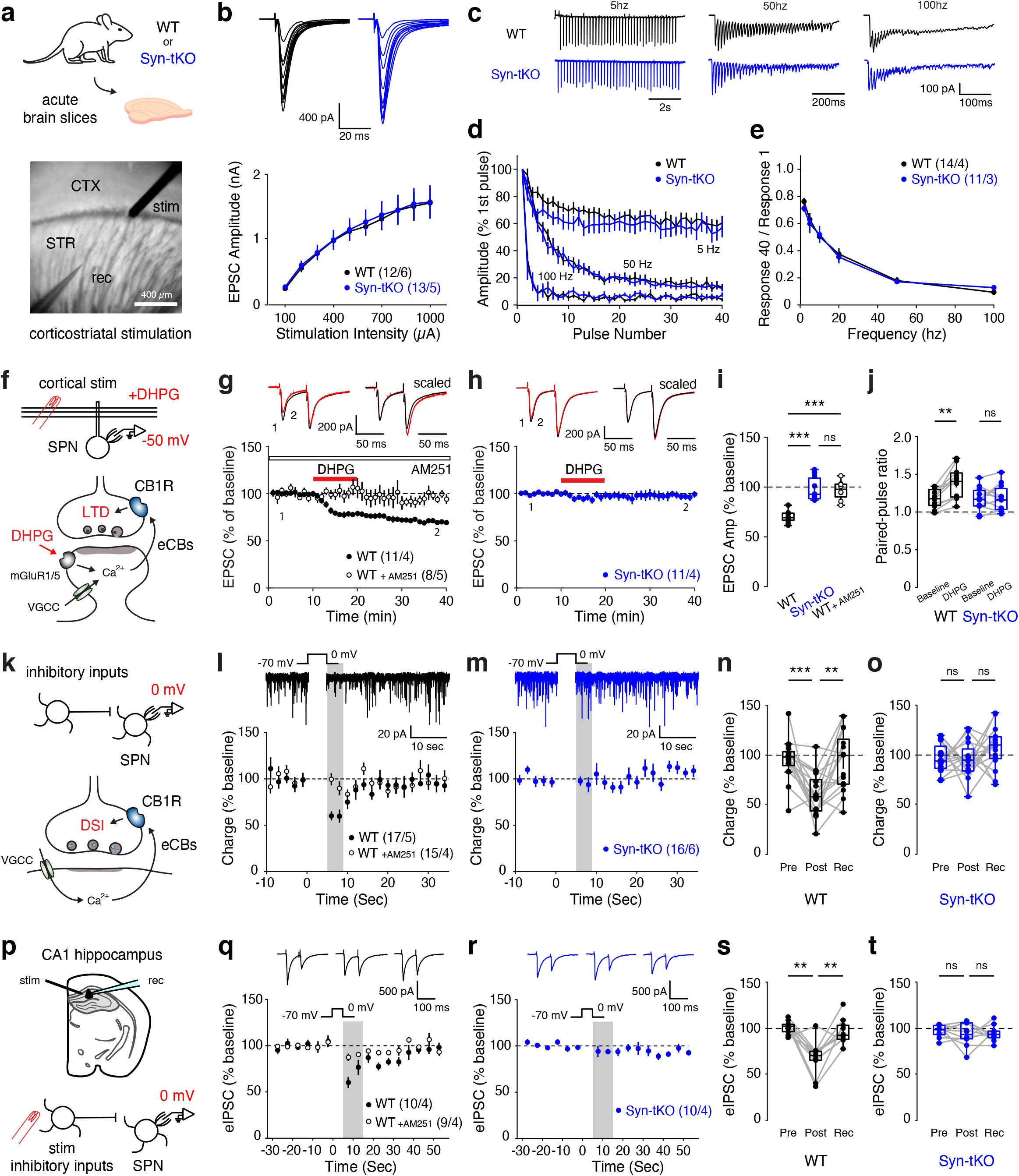
Endocannabinoid-dependent LTD and DSI are impaired in Syn-tKO mice. **a**, Acute slices of dorsal striatum prepared for whole-cell recordings and stimulation of corticostriatal transmission. **b**, Top, representative traces of evoked corticostriatal EPSCs across a range of stimulation intensities. Bottom, normal input-output curves in Syn-tKO mice (WT: n = 12 cells / 6 mice; Syn-tKO: n = 13 cells / 6 mice; p = 0.888). **c**, Representative traces of responses to repeated stimulation across a range of stimulation frequencies. **d, e**, No difference in use-dependent synaptic properties in Syn-tKO mice, as measured by short-term depression dynamics (**d**; WT: n = 14 cells / 4 mice; Syn-tKO: n = 11 cells / 3 mice; 5 Hz: p = 0.304; 50 Hz: p = 0.651; 100 Hz: p = 0.691) and steady-state amplitudes (**e**; p = 0.756) in response to repeated stimulation across a range of frequencies. **f**, eCB-LTD experimental strategy. Top, schematic of whole-cell recordings from SPNs during induction of eCB-LTD; bottom, eCB-LTD induction pathway engaged with DHPG (50 µM) and depolarization (−50 mV). **g-i**, Summary of DHPG-mediated eCB-LTD in WT mice (**g**), which is fully blocked by the CB1R antagonist AM251 (10 µM) (WT: n = 11 cells / 4 mice, 69.95 ± 1.70%; WT + AM251: n = 8 cells / 5 mice, 96.89 ± 3.42%; p = 6.375e-4); top, representative traces. (**h**) eCB-LTD is impaired in Syn-tKO mice compared to WT (Syn-tKO: n = 11 cells / 4 mice, 97.76 ± 3.49%; p = 2.106e-4); top, representative trace. **j**, Significant increase in WT PPRs (baseline: 1.18 ± 0.04; post-DHPG: 1.39 ± 0.06; p = 0.001) but not in Syn-tKO PPRs (baseline: 1.17 ± 0.05; post-DHPG: 1.19 ± 0.05; p = 0.831). **k**, DSI experimental strategy. Top, schematic of whole-cell recordings from SPNs during induction of DSI; bottom, DSI induction pathway engaged with strong depolarization (0 mV). **l**, Summary of DSI in WT mice, which is blocked by AM251 (10 µM); top, representative WT experiment. **m**, DSI is impaired in Syn-tKO mice; top, representative Syn-tKO experiment. **n**, DSI summary for WT mice (n = 17 cells / 5 mice, pre-depol: 95.64 ± 5.01%, post-depol: 59.91 ± 5.28%, recovery: 93.51 ± 7.20%, p = 5e-4, p = 1.6e-3). **o**, DSI summary for Syn-tKO mice (n = 16 cells / 6 mice, pre-depol: 97.25 ± 3.66%, post-depol: 95.70 ± 4.54%, recovery: 107.59 ± 5.20%, p = 0.959, p = 0.148). **p**, Schematic of evoked DSI experiments in CA1 of the hippocampus. **q, s**, Summary of DSI in recorded principal neurons in CA1 of WT mice (n = 10 cells / 4 mice; pre-depol: 101.50 ± 2.18%; post-depol: 68.26 ± 5.85%; recovery: 96.77 ± 4.36%; p = 3.9e-3, p = 3.9e-3). **r, t**, Hippocampal DSI is impaired in Syn-tKO (n = 10 cells / 4 mice; pre-depol: 97.24 ± 2.08%; post-depol: 93.70 ± 3.94%; recovery: 94.73 ± 2.72%; p = 0.625, p = 0.846). Data are mean ± s.e.m. Statistical significance was assessed by 2-way repeated measures ANOVA with multiple comparisons (**b, d, e**), ANOVA with multiple comparisons (**i**), and by Wilcoxon signed tests (**j, n, o, s, t**) (*** p < 0.001; ** p < 0.01; n.s. non-significant).

We next measured the use-dependent dynamics of synaptic transmission by delivering stimulus trains at varying frequencies (Fig. 1c). We measured the rate of synaptic depression resulting from repeated stimulation^23^ and found virtually indistinguishable depression dynamics (Fig. 1d) and steady-state response amplitudes (Fig. 1e) between WT and Syn-tKO cells across stimulation frequencies. Together, these results show that basal corticostriatal synaptic transmission in Syn-tKO mice is largely normal, including responses engaged by repeated stimuli that depend on the rates of presynaptic vesicle recycling and the sizes of the reserve vesicle pool.

## Syn-tKO mice lack eCB-dependent plasticity

One of the best-characterized forms of corticostriatal synaptic plasticity is eCB-LTD^43,44,45^ that is required for striatal learning^33,34^. Importantly, impairments in corticostriatal eCB-LTD are observed in mouse models of PD^32,46,47^. We assayed eCB-LTD in acute slices of young-adult (3 months old) WT and Syn-tKO mice by combining slight membrane depolarization (−50 mV) with an application of a type I mGluR agonist ((S)-3,5-dihydroxyphenylglycine (DHPG; 50 µM); Fig. 1f), which results in a lasting depression of evoked corticostriatal excitatory postsynaptic currents (EPSCs) (Fig. 1g). Strikingly, we found that eCB-LTD is abolished in Syn-tKO mice (Fig. 1h). Syn-tKO cells were indistinguishable from WT cells in the presence of the CB1R antagonist, AM251 (10 µM) (Fig. 1g,i). Importantly, paired-pulse ratios (PPRs) were significantly increased in WT cells following eCB-LTD, but not in Syn-tKO cells (Fig. 1j), consistent with a selective decrease in presynaptic release probability in WT cells. We observed impaired eCB-LTD in both young-adult and aged mice (16-18 months old; Extended Data Fig. 1c-f), suggesting that the phenotype is not an age-dependent effect, but instead due to a direct loss of an endogenous synuclein function. Furthermore, we found that eCB-LTD was normally expressed in KO mice lacking α-Syn alone or both β- and γ-synuclein (βγ-Syn-KO mice), suggesting redundancy among synucleins (Extended Data Fig. 2a-d).

In order to further characterize the Syn-tKO phenotype, we measured depolarization-induced suppression of inhibition (DSI)^38^, a different form of eCB-dependent plasticity in the striatum. During DSI, strong depolarization of SPNs results in the Ca^2+^-dependent synthesis and release of eCBs that transiently suppress inhibitory inputs (Fig. 1k)^35,36,37,48^. Indeed, a 5-second depolarization (to 0 mV) in WT cells was sufficient to transiently inhibit spontaneous inhibitory postsynaptic currents (sIPSCs) in a CB1R-dependent manner (Fig. 1l,n). Strikingly, the same DSI protocol failed to elicit a significant reduction in sIPSCs in Syn-tKO mice (Fig. 1m,o). We observed the same results when we repeated this experiment using a stimulation-evoked IPSC protocol (Extended Data Fig. 3a-d), with WT but not Syn-tKO cells showing a significant increase in PPRs during DSI (Extended Data Fig. 3e), which reflects the presynaptic locus of the transient suppression of inhibitory inputs.

Finally, in a parallel set of experiments, we recorded DSI in pyramidal neurons of the hippocampal CA1 region (Fig. 1p)^37^. Here we once again found that DSI was readily inducible in WT cells, but not in Syn-tKO cells (Fig. 1q-t; Extended Data Fig. 3f-h). The observations that Syn-tKO mice exhibit impairments in two forms of eCB plasticity (eCB-LTD and DSI), across different synapse types (glutamatergic and GABAergic), and brain regions (striatum and hippocampus) suggest a broad defect in eCB signaling in Syn-tKO mice.

## Presynaptic CB1Rs are intact in Syn-tKO mice

α-Syn is thought to function predominantly in the presynaptic terminal, suggesting that the impairment in eCB-dependent synaptic plasticity in Syn-tKO mice is likely due to a failure of CB1R signaling^49^. To test this hypothesis, we applied the CB1R agonist WIN55,212 (WIN; 2 µM) in acute brain slices. WIN strongly depressed corticostriatal transmission via direct activation of presynaptic CB1Rs, bypassing the postsynaptic eCB synthesis and release mechanisms engaged during eCB-LTD and DSI (Fig. 2a). We found that WIN strongly reduced evoked EPSCs in both WT and Syn-tKO mice (Fig. 2b,c). The magnitude of synaptic depression was indistinguishable between genotypes (Fig. 2d), as was the concomitant significant increase in PPRs (Fig. 2e) that would be expected for a presynaptic weakening via CB1R activation. These results were reproduced when repeated in aged mice (Extended Data Fig. 4a-d). Thus, presynaptic CB1R function is intact in Syn-tKO mice, suggesting a postsynaptic deficit upstream of CB1R activation.

**Figure 2.**
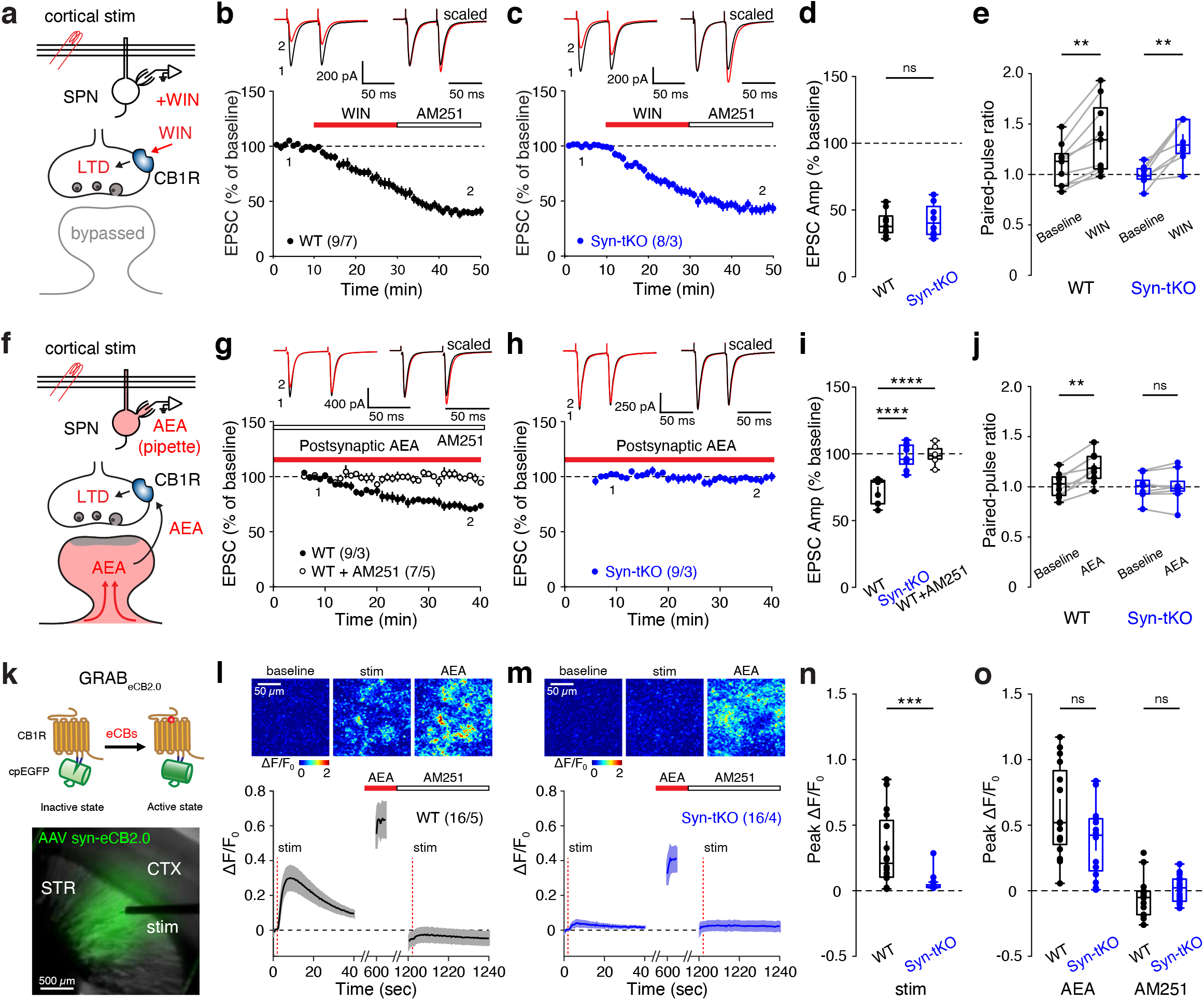
eCB release is impaired in Syn-tKO mice. **a**, WIN-LTD experimental strategy. Top, schematic of whole-cell recordings from SPNs during induction of presynaptic LTD via WIN (2 µM) application; bottom, WIN-mediated LTD bypasses postsynaptic eCB synthesis and release steps of eCB-LTD. **b-d**, WIN application results in significant corticostriatal LTD in both WT and Syn-tKO mice (WT: n = 9 cells / 7 mice, 39.91 ± 3.18%; Syn-tKO: n = 8 cells / 3 mice, 42.46 ± 4.36%; p = 0.673). **e**, Significant increases in PPRs in both WT (baseline: 1.09 ± 0.07; post-WIN: 1.36 ± 0.12; p = 3.9e-3) and Syn-tKO mice (baseline: 0.99 ± 0.04; post-WIN: 1.33 ± 0.07; p = 7.8e-3). **f**, Schematic of AEA-loading experiments. Top, whole-cell recordings from SPNs while dialyzing cells with AEA (50 µM) through the patch-pipette; bottom, AEA-loading bypasses postsynaptic eCB synthesis steps of eCB-LTD. **g-i**, AEA-loading results in progressive LTD in WT cells, which is blocked in the presence of AM251 (10 µM) (WT: n = 9 cells / 3 mice, 72.40 ± 3.10%; WT + AM251: n = 7 cells / 5 mice, 99.13 ± 2.63 %; p = 1.08e-5), but AEA-loading does not result in significant LTD in Syn-tKO cells (Syn-tKO: n = 9 cells / 3 mice, 98.16 ± 3.05; p = 7.08e-6). **j**, Significant PPR increase in WT (baseline: 1.02 ± 0.04; end: 1.19 ± 0.05; p = 3.9e-3) but not Syn-tKO cells (baseline: 1.00 ± 0.04; end: 1.00 ± 0.05; p = 0.82). **k**, Schematic of GRAB_eCB2.0_ experiments. Top, cartoon of GRAB_eCB2.0_ activation in the presence of eCBs; bottom, acute slice expressing GRAB_eCB2.0_ in dorsal striatum. **l**, Electrical stimulation resulted in transient increases in GRAB_eCB2.0_ signal in WT mice; top, representative images. **m, n**, Compared to WT slices, stimulation-evoked GRAB_eCB2.0_ transients in Syn-tKO slices were significantly reduced (WT: n = 16 slices / 5 mice, 0.31 ± 0.07 ΔF/F_0_; Syn-tKO: n = 16 slices / 4 mice, 0.05 ± 0.02 ΔF/F_0_; p = 8.52e-4). **l, m, o**, Following stimulation experiments, all slices were subsequently bath applied with AEA (50 µM, 10 minutes) and then AM251 (10 µM, 10 minutes) as positive and negative controls, respectively, (**o**) with no differences observed between GRAB_eCB2.0_ signals of WT and Syn-tKO slices in the presence of AEA (WT: 0.61 ± 0.09 ΔF/F_0_; Syn-tKO: 0.37 ± 0.07 ΔF/F_0_; p = 0.073) or AM251 (WT: -0.05 ± 0.04 ΔF/F_0_; Syn-tKO: 0.02 ± 0.03 ΔF/F_0_; p = 0.086). Data are mean ± s.e.m. Statistical significance was assessed by Mann-Whitney tests (**d, n, o**), Wilcoxon signed tests (**e, j**), and ANOVA with multiple comparisons (**i**) (**** p < 0.0001; *** p < 0.001; ** p < 0.01; n.s. non-significant).

## Release of eCBs is impaired in Syn-tKO mice

Given the defects in eCB plasticity across different experimental contexts, we next tested whether a more upstream step in eCB signaling was impaired in Syn-tKO mice, namely the postsynaptic release of eCBs as retrograde signals. Postsynaptic release of eCBs precedes CB1R activation but is downstream of eCB synthesis^40,50^. Although the specific mechanisms of retrograde eCB release are not well understood^50,51^, the direct introduction of eCBs into a postsynaptic neuron via a patch pipette has been shown to induce a progressive release of these eCBs, resulting in synaptic depression^52,53^. Thus, in order to directly test eCB release, we dialyzed SPNs intracellularly with the endogenous eCB anandamide (AEA; 50 µM) through the patch-pipette (Fig. 2f). In WT cells, the intracellular application of AEA caused a progressive depression of evoked corticostriatal EPSCs that depended on CB1R function (Fig. 2g). Strikingly, in Syn-tKO cells postsynaptic AEA loading had no effect (Fig. 2h,i). Correspondingly, we observed significant PPR increases in WT cells, but not in Syn-tKO cells (Fig. 2j). Because intracellular loading with AEA bypasses the eCB synthesis pathways, these results suggest that the defect in Syn-tKO mice lies specifically in the release of eCBs from postsynaptic cells.

To directly visualize eCB release, we utilized a recently developed eCB fluorescent sensor (GRAB_eCB2.0_)^54^. Viral expression of the GRAB_eCB2.0_ sensor in the dorsal striatum of mice allowed us to image stimulation-induced release of eCBs in acute slices (Fig. 2k). Local electrical stimulation in WT slices resulted in a significant increase in GRAB_eCB2.0_ signal, reflecting the release of eCBs (Fig. 2l). However, evoked GRAB_eCB2.0_ signals were significantly reduced in Syn-tKO mice (Fig. 2m,n), consistent with a deficit in eCB release. Importantly, we validated GRAB_eCB2.0_ sensor expression and function in all imaged slices. Bath application of AEA (10 µM) significantly increased GRAB_eCB2.0_ fluorescence in both WT and Syn-tKO mice, and AM251 (10 µM) decreased GRAB_eCB2.0_ fluorescence and blocked stimulation-induced GRAB_eCB2.0_ activity in WT slices (Fig. 2l-o). Thus, in combination with our electrophysiology data, these results suggest that normal eCB release requires synucleins.

## eCB plasticity requires postsynaptic α-Syn

Thus far, our results suggest that synucleins are required for the postsynaptic release of eCBs. To further test this conclusion, we sparsely infected SPNs in the dorsolateral striatum of Syn-tKO mice with adeno-associated viruses (AAVs) that co-express GFP and α-Syn (Fig. 3a, top). Recordings of corticostriatal eCB-LTD from GFP+ or GFP-cells allowed us to directly test whether postsynaptic exogenous α-Syn can rescue the Syn-tKO phenotype (Fig. 3a, bottom). As expected, eCB-LTD was not observed in GFP-cells (Fig. 3b). Remarkably, almost all GFP+ cells expressing α-Syn exhibited significant eCB-LTD (10 out of 11) (Fig. 3c,e). The presence or absence of α-Syn in recorded cells was confirmed by immunocytochemistry (Fig. 3b,c, top). Moreover, viral expression of C-terminally truncated α-Syn (residues 1-95) also rescued eCB-LTD in Syn-tKO cells (Fig. 3d,e). The rescued eCB-LTD in GFP+ cells was accompanied by a significant increase in PPRs, which was not observed in uninfected GFP-cells (Fig. 3f). Finally, postsynaptic rescue of α-Syn also restored striatal DSI in Syn-tKO cells (Fig. 3h,i). Together, these results demonstrate that not only are synucleins required for eCB plasticity, but also that the role they play is a postsynaptic one.

**Figure 3.**
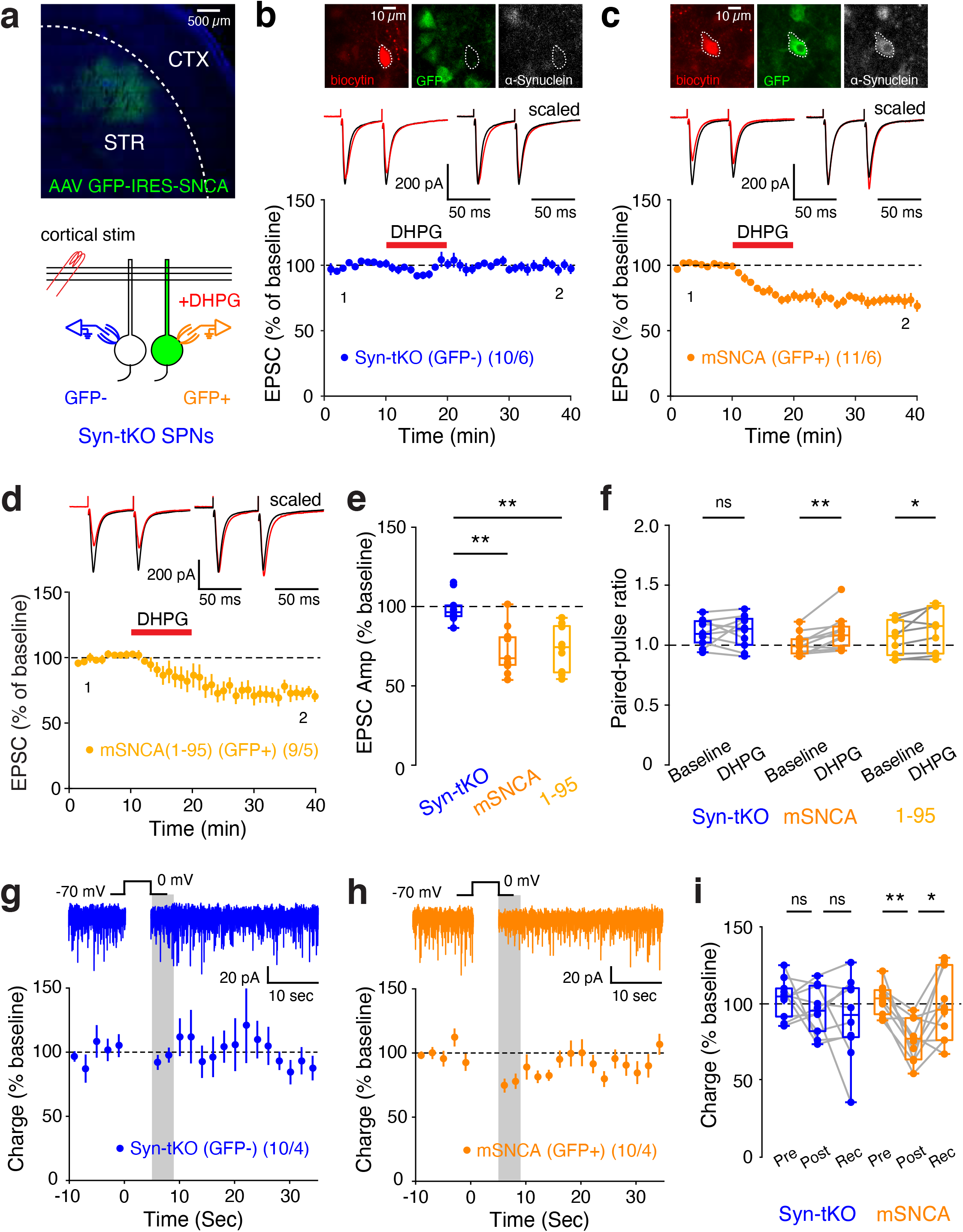
Postsynaptic α-Syn rescues eCB plasticity in Syn-tKO mice. **a**, Experimental approach for eCB-LTD recordings following postsynaptic viral rescue of α-Syn in Syn-tKO mice. Top, AAV-mediated expression of α-Syn and GFP in dorsolateral striatum of Syn-tKO mice; bottom, Syn-tKO (GFP-) and α-Syn-expressing SPNs (GFP+) targeted for recordings. **b, c**, Top, Post-hoc staining of biocytin-filled recorded cells for GFP and α-Syn expression. **b, c, e**, Viral postsynaptic delivery of α-Syn into Syn-tKO cells is sufficient to rescue eCB-LTD (GFP-, pooled: n = 10 cells / 6 mice, 99. 04 ± 2.89%; GFP+, mSNCA: n = 11 cells / 6 mice, 72.71 ± 4.32%; p = 2.09e-4). (**d, e**) Viral expression of a full C-terminus truncated α-Syn (1-95) still rescued eCB-LTD in Syn-tKO mice (GFP+, 1-95: n = 9 cells / 5 mice, 73.47 ± 4.84%; p = 5.41e-4). **f**, Significant increases to PPR in α-Syn-expressing cells (GFP+, mSNCA; baseline: 1.01 ± 0.03; post-DHPG: 1.11 ± 0.04; p = 2.0e-3) and C-terminus truncated α-Syn-expressing cells (GFP+, 1-95; baseline: 1.07 ± 0.05; post-DHPG: 1.14 ± 0.06; p = 0.039), but not GFP-cells (GFP-, pooled; baseline: 1.08 ± 0.03; post-DHPG: 1.09 ± 0.04; p = 1.0). **g-i**, DSI remains absent in Syn-tKO SPNs uninfected with α-Syn (GFP-cells: n = 10 cells / 4 mice, pre-depol: 103.58 ± 4.01%, post-depol: 95.81 ± 4.88%, recovery: 90.66 ± 8.44%, p = 0.695, p = 0.770), but is rescued in infected cells (GFP+ cells: n = 10 cells / 4 mice, pre-depol: 102.73 ± 3.19%, post-depol: 76.21 ± 4.29%, recovery: 98.32 ± 7.27%, p = 3.9e-3, p = 0.027). Data are mean ± s.e.m. Statistical significance was assessed by Wilcoxon signed tests (**f, i**), and ANOVA with multiple comparisons (**e**) (** p < 0.01; * p < 0.05; n.s. non-significant).

## Membrane-binding domains of α-Syn are required for eCB plasticity

In order to dissect the mechanism of synuclein function in postsynaptic eCB release, we sparsely expressed α-Syn in the striatum of Syn-tKO mice as before, but included mutations in the α-Syn rescue sequence to determine which regions (and therefore functions) of α-Syn are required for eCB-dependent plasticity. Although we previously observed that C-terminal truncation of α-Syn had no effect on eCB-LTD (Fig. 3d), we asked if C-terminal serine 129, a site previously implicated in Ca^2+^-binding affinity and regulating PD neurodegeneration^55^, could modulate eCB-LTD. However, we found that phosphorylation at serine 129 was not relevant for α-Syn’s function within eCB-LTD, as neither alanine (S129A, phosphorylation-deficient) nor aspartate substitutions (S129D, phosphorylation-mimic)^56^ affected the viral rescue of eCB-LTD in Syn-tKO mice (Extended Data Fig. 5).

Our results thus indicate that the N-terminal domain of α-Syn is required for eCB release. The major biochemical activity of α-Syn consists of phospholipid membrane binding that is mediated by its N-terminal domain^13,15,57^. To test whether membrane binding by α-Syn is required for eCB-LTD, we virally expressed α-Syn mutants carrying A11P and V70P (A11P/V70P) substitutions that ablate membrane binding by α-Syn but do not impair its synaptic localization^15^. Remarkably, A11P/V70P-mutant α-Syn failed to rescue eCB-LTD in Syn-tKO mice (Fig. 4a-c), suggesting that membrane binding of α-Syn is required for eCB-LTD. To strengthen this hypothesis, we repeated these experiments in cells infected with A30P mutant α-Syn, a PD mutation that also decreases lipid binding by α-Syn^15^. A30P-mutant α-Syn also did not rescue the loss of eCB-LTD in Syn-tKO mice (Fig. 3d,e). Correspondingly, PPRs were increased in cells expressing WT α-Syn but not in cells expressing A11P/V70P- or A30P-mutant α-Syn (Fig. 3f). Together, these results demonstrate that in postsynaptic neurons, α-Syn enables eCB-LTD by binding to phospholipid membranes, likely by mediating the postsynaptic release of eCBs.

**Figure 4.**
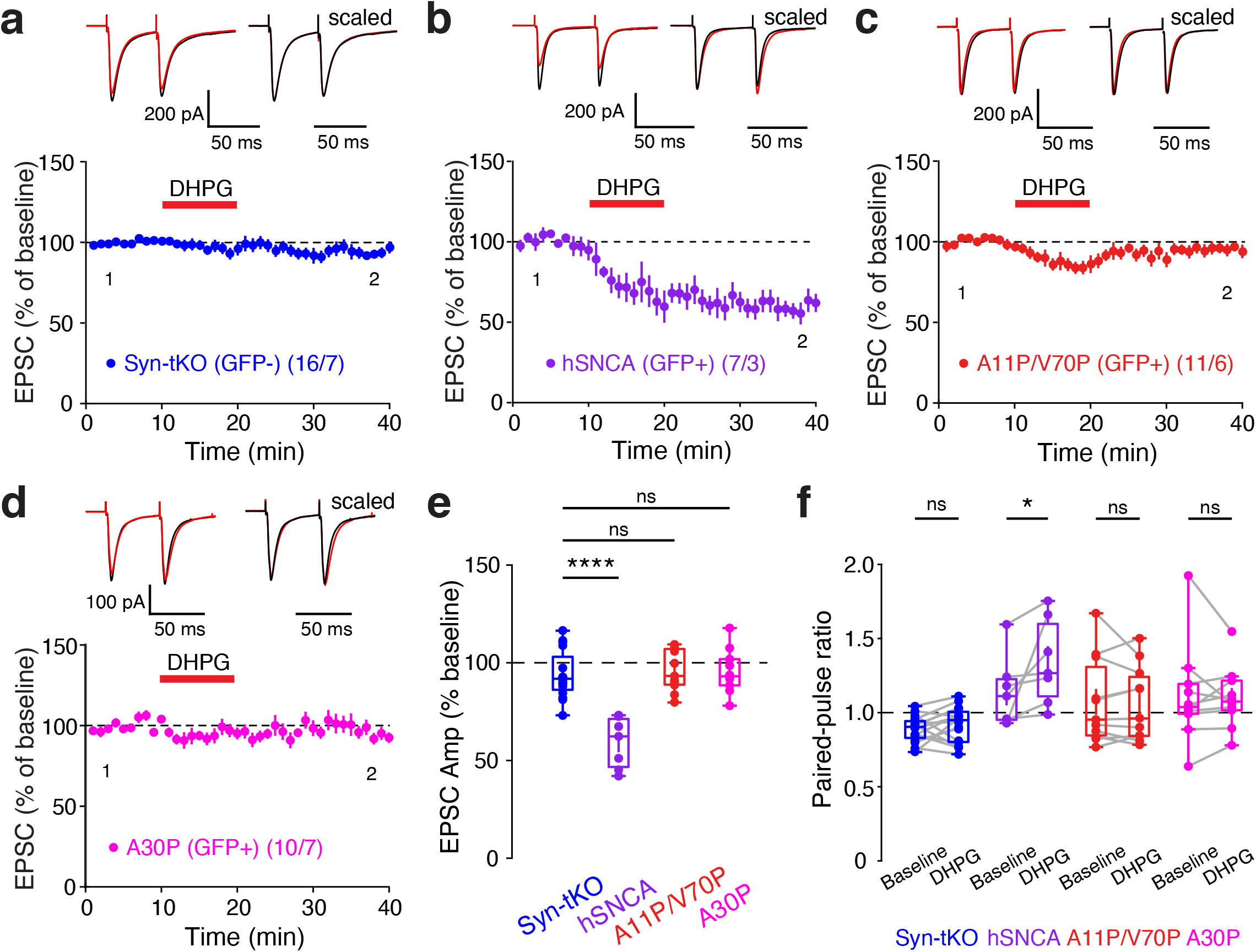
Postsynaptic membrane interaction by α-Syn is required for eCB-LTD. **a-e**, eCB-LTD could not be rescued by the expression of (**c**) mutant A11P/V70P α-Syn (GFP-, pooled: n = 16 cells / 7 mice, 93.84 ± 3.06%; GFP+, hSNCA: n = 7 cells / 3 mice, 59.08 ± 4.90%, p = 4.98e-7; A11P/V70P: n = 11 cells / 6 mice, 95.40 ± 3.17%; p = 0.986) or by (**d**) A30P α-Syn (GFP+, A30P: n = 10 cells / 7 mice, 95.75 ± 3.59%; p = 0.977). **f**, Significant increase in PPRs for Syn-tKO cells infected with α-Syn (GFP+, hSNCA; baseline: 1.14 ± 0.09; post-DHPG: 1.35 ± 0.11; p = 0.016), but no significant increase in PPRs observed in uninfected cells (GFP-, pooled; baseline: 0.89 ± 0.02; post-DHPG: 0.92 ± 0.03; p = 0.255) or in cells infected with A11P/V70P (baseline: 1.07 ± 0.09; post-DHPG: 1.05 ± 0.08; p = 0.496) or A30P mutant α-Syn (baseline: 1.12 ± 0.11; post-DHPG: 1.10 ± 0.07; p = 0.695). Data are mean ± s.e.m. Statistical significance was assessed by ANOVA with multiple comparisons (**e**) and Wilcoxon signed test (**f**) (**** p < 0.0001; * p < 0.05; n.s. non-significant).

## Postsynaptic SNAREs are required for eCB release

α-Syn has been shown to act as a SNARE chaperone that facilitates SNARE complex assembly during vesicular exocytosis by binding to phospholipid membranes^4,24,58^. SNARE proteins not only mediate presynaptic vesicle exocytosis but are also essential for postsynaptic exocytosis of AMPA receptors and other proteins^59,60,61^. Thus, the fact that eCB release requires postsynaptic α-Syn that is competent to bind to phospholipid membranes suggests that eCBs are released by synuclein-dependent exocytosis. To investigate this possibility, we tested if postsynaptic SNAREs are involved in eCB-dependent plasticity and eCB release.

We sparsely infected SPNs in the dorsolateral striatum of WT mice with lentiviruses that co-express GFP and tetanus-toxin light chain (TeNT), which inactivates synaptobrevin-2, a SNARE protein involved in most forms of exocytosis. We confirmed that postsynaptic TeNT expression did not disrupt basal synaptic properties of infected SPNs, as previously shown for hippocampal neurons^62,63^ (Extended Data Fig. 6). Next, we measured eCB-dependent plasticity, comparing GFP+ (TeNT-expressing) cells to adjacent GFP-controls. Strikingly, TeNT significantly impaired eCB-LTD (Fig. 5a-c) and blocked DSI (Fig. 5d-f), an effect that was not revealed in previous studies using acute neurotoxin dialysis through the patch-clamp recording pipette^37^. Together, the impaired eCB-LTD and DSI results mirror the Syn-tKO phenotypes and suggest that postsynaptic SNAREs are also required for eCB-dependent plasticity. Lastly, to further explore the specificity of the effect of TeNT in impairing the release of eCBs, we performed the AEA-loading experiment as before. AEA-loading of GFP+ cells expressing TeNT failed to induce progressive synaptic depression, whereas loading of GFP-control cells robustly suppressed synaptic transmission (Fig. 5g-i). Thus, in addition to synucleins, SNAREs are required postsynaptically for the active release of eCBs (Fig. 5j).

**Figure 5.**
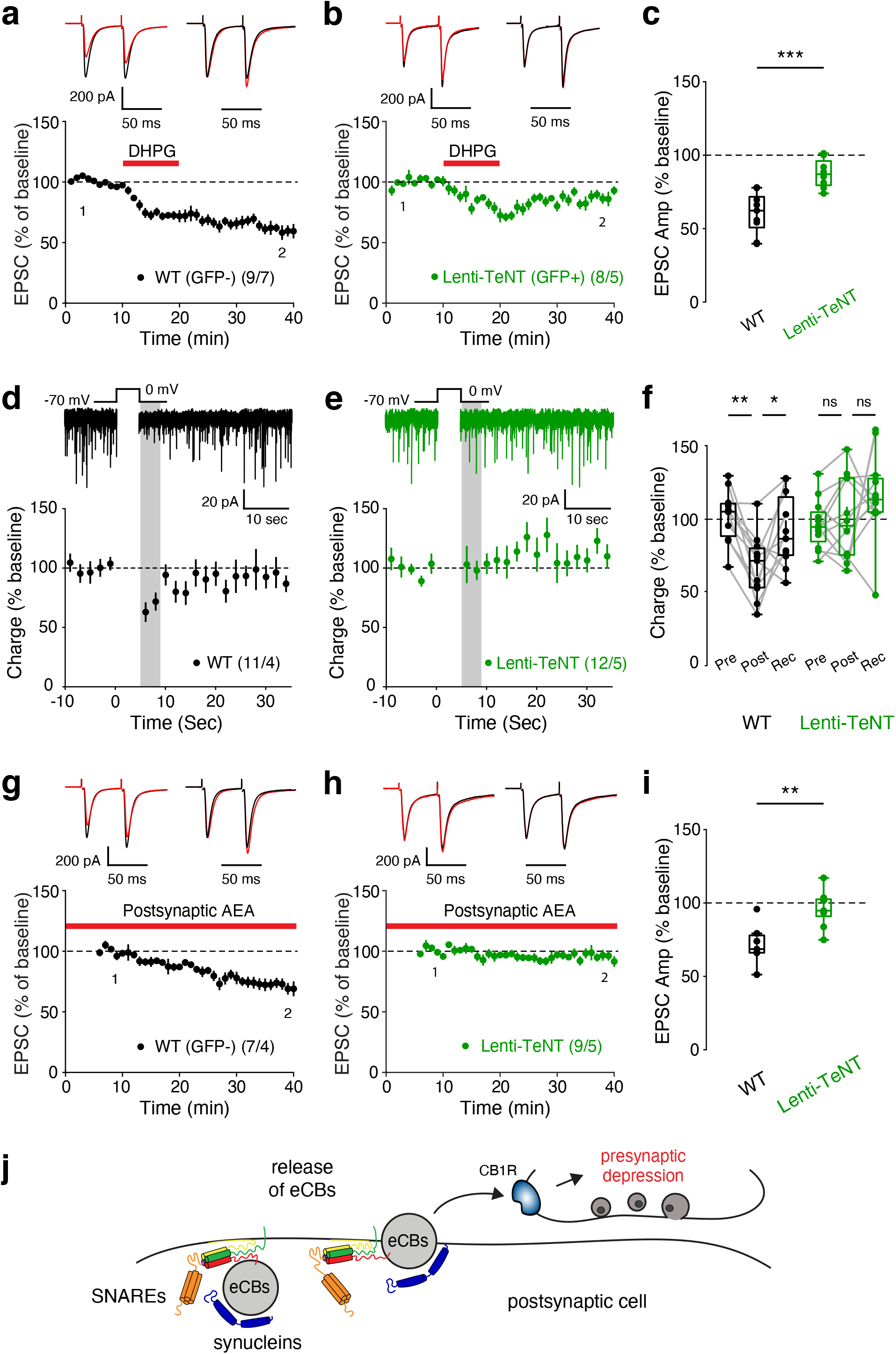
Postsynaptic SNAREs are required for normal eCB plasticity and eCB release. **a-c**, Postsynaptic lentiviral expression of TeNT impairs eCB-LTD (GFP-: n = 9 cells / 7 mice, 60.47 ± 4.73; GFP+: n = 8 cells / 5 mice, 87.54 ± 3.51%; p = 9.87e-4). **d-f**, Postsynaptic TeNT significantly impairs DSI (GFP-: n = 11 cells / 4 mice, pre-depol: 101.93 ± 5.38%, post-depol: 67.33 ± 6.50%, recovery: 91.48 ± 7.40%, p = 9.77e-4, p = 0.032; GFP+: n = 12 cells / 5 mice, pre-depol: 97.40 ± 4.24%, post-depol: 99.58 ± 7.00%, recovery: 111.00 ± 6.81%, p = 0.569, p = 0.339). **g-i**, Postsynaptic TeNT prevents AEA-loading LTD (GFP-: n = 7 cells / 4 mice, 71.57 ± 5.20%; GFP+: n = 9 cells / 5 mice, 96.06 ± 4.05%, p = 5.2e-3). **j**, Hypothetical model depicting postsynaptic synucleins and SNAREs coordinating the release of eCBs. Data are mean ± s.e.m. Statistical significance was assessed by Mann-Whitney tests (**c, i**) and Wilcoxon signed tests (**f**) (*** p < 0.001; ** p < 0.01; * p < 0.05; n.s. non-significant).

## Summary

Here we show that eCBs are released by postsynaptic vesicular exocytosis, in a process that requires synucleins. Thus, we report an unexpected convergence of two puzzling questions in neuroscience, namely the questions of the function of synucleins and of the mechanism of eCB release. We show that mice lacking all three synuclein isoforms have apparently normal basal synaptic properties but exhibit significant defects in multiple forms of eCB-dependent plasticity spanning different time frames (eCB-LTD and DSI), synapse types (glutamatergic and GABAergic), and brain regions (striatum and hippocampus). Using direct measurements of eCB release, we demonstrate that synuclein-deficient neurons suffer from a loss of eCB release, but retain normal CB1R function. Strikingly, bypassing the Ca^2+^-dependent eCB synthesis processes via postsynaptic loading of neurons with AEA, an endogenous eCB, revealed that the export of AEA from the postsynaptic cell is impaired by the synuclein deletion. Mechanistically, we identify the N-terminal membrane-binding domain of α-Syn, as well as postsynaptic synaptobrevin SNAREs, as required for eCB release. Together, these results point towards vesicular exocytosis as the process underlying eCB transmission.

Our results are surprising given that α-Syn is known to function presynaptically, and do not preclude additional presynaptic roles for α-Syn. Our viral α-Syn rescue experiments take advantage of the corticostriatal circuit’s compartmentalization of pre- and postsynaptic cells to demonstrate that the postsynaptic expression of α-Syn is sufficient to restore eCB-dependent plasticity in Syn-tKO mice. Indeed, synuclein-dependent release of eCBs adds to our growing understanding of the roles played by postsynaptic SNAREs during synaptic plasticity^60,61^. Furthermore, our results provide a potential link between eCB signaling and PD. In particular, our finding that A30P PD mutant α-Syn is unable to rescue eCB-LTD suggests that eCB release and eCB-dependent plasticity may be aberrant in PD, potentially contributing to the cognitive deficits observed in PD pathology. Additionally, our results are unexpected since an early report suggested that SNAREs are not involved in eCB release^37^. However, the previous study relied on acute Botulinum toxin light-chain dialysis, which may be temporally insufficient to fully access SNAREs and block eCB-release. In our study, we achieved neurotoxin expression (e.g., Lenti-GFP-TeNT) that enables TeNT action for multiple days prior to experiments. Together, our results demonstrate a novel postsynaptic function of endogenous synucleins in regulating eCB release and synaptic plasticity, and reveal that eCBs are released postsynaptically via synuclein-dependent vesicular exocytosis.

## Acknowledgments

This study was supported by grants from the NINDS/NIH P50NS091857 (T.C.S), R01NS091144 (J.B.D.), U01NS113358 (J.B.D.), HHMI Gilliam Fellowship (E.A.), NSF GRFP (E.A.), and Stanford DARE Fellowship (E.A.). We thank Xiaobai Ren for technical support and the members of Ding lab for valuable discussions.

## Author contributions

E.A., T.C.S, and J.B.D. conceptualized the project and designed the experiments. E.A. performed electrophysiology and imaging experiments. Y.S., Y.L, K.R., and S.W generated α-Syn and TeNT viral tools. A.D. and Y.Li. generated the GRAB_eCB2.0_ sensor. E.A., T.C.S., and J.B.D. analyzed the data. E.A., T.C.S., and J.B.D. wrote the manuscript with input from all authors.

## LEAD CONTACT AND MATERIALS AVAILABILITY

This study did not generate new unique reagents. Further information and requests for resources and reagents should be directed to and will be fulfilled by the Lead Contacts, Thomas C. Südhof and Jun B. Ding.

## EXPERIMENTAL MODEL AND SUBJECT DETAILS

### Animals

All experiments were performed in accordance with protocols approved by the Stanford University Animal Care and Use Committee in keeping with the National Institutes of Health’s *Guide for the Care and Use of Laboratory Animals*. Both male and female mice were used for all experiments at ∼3-months old (P70-P100), with the exception of recordings from aged mice (16-18 months old). Syn-tKO mice (α-Syn^-/-^;β-Syn^-/-^;γ-Syn^-/-^) were generated as previously described^42^. WT C57BL/6 mice were maintained as controls, and Syn-tKO mice were back-crossed to C57BL/6 every 6-10 months in order to maintain a consistent background between Syn-tKO and WT lines. α-Syn-KO (α-Syn^-/-^) and βγ-Syn-KO (β-Syn^-/-^; γ-Syn^-/-^) were generated from these backcrosses. Stereotaxic injections were performed 2-6 weeks before recordings.

## METHODS DETAILS

### Acute brain slice preparation

Adult mice (male and female) were anesthetized with isoflurane, decapitated, and brains were extracted and briefly submerged into chilled artificial cerebrospinal fluid (ACSF) containing 125 mM NaCl, 2.5 mM KCl, 1.25 mM NaH_2_PO_4_, 25 mM NaHCO_3_, 15 mM glucose, 2 mM CaCl_2_, and 1 mM MgCl_2_, oxygenated with 95% O_2_ and 5% CO_2_ (300-305 mOsm, pH 7.4). Oblique horizontal slices (300 µm thickness) containing dorsal striatum (or coronal slices containing hippocampus) were then prepared using a tissue vibratome (VT1200S, Leica), incubated in chambers containing 34°C ACSF for 30 min, and then allowed to recover at room temperature for 30 min. After recovery, slices were transferred to a submerged recording chamber perfused with ACSF at a rate of 2-3 ml/min at a temperature of 30-31°C. All recordings were performed within 5hrs of slice recovery.

### Whole-cell slice electrophysiology

Whole-cell voltage clamp recordings were made with glass pipettes (3-4 MΩ) filled with internal solution containing 126 mM CsMeSO_3_, 10 mM HEPES, 1 mM EGTA, 2 mM QX-314 chloride, 0.1 mM CaCl_2_, 4 mM Mg-ATP, 0.3 mM Na_3_-GTP, and 8 mM disodium phosphocreatine (280-290 mOsm, pH 7.3 with CsOH), and cells were voltage clamped at -70 mV unless specified otherwise. Access resistance was measured by injection of hyperpolarizing pulses (−5 mV, 100 µs) and was less than 25 MΩ for all recordings and only cells with a change in access resistance <20% throughout the entire experiment were included in the analysis. Similarly, input resistance was monitored throughout the entirety of experimental recordings. For EPSC recordings, 50 µM Picrotoxin was added to block GABA_A_ receptor-mediated currents. Evoked EPSCs were elicited by stimulating axons via a concentric bipolar stimulating electrode (FHC). Whole-cell patch clamp recordings were performed using a Multiclamp 700B (Molecular Devices), monitored with WinWCP (Strathclyde Electrophysiology Software) and analyzed offline using Clampfit 10.0 (Molecular Devices) and custom-made MATLAB (Mathworks) software. Signals were filtered at 2 kHz and digitized at 10 kHz (NI PCIe-6259, National Instruments).

### Basal corticostriatal synaptic activity recordings

For input-output curves of corticostriatal synapses, 3 EPSCs were averaged at stimulation intensities ranging from 100 µA to 1000 µA (100 µA step size) and the average amplitude measured. For measuring dynamics of repeated stimulation, trains of 40 stimulation pulses were delivered at a range of frequencies (2, 5, 10, 20, 50, 100 Hz)^23^. Miniature excitatory postsynaptic currents (mEPSCs) were measured by continuously recording for 10 min in the presence of 1 µM Tetrodotoxin to prevent action potential firing and 50 µM Picrotoxin to block GABA_A_ receptor-mediated currents.

### eCB-LTD recordings

For long-term eCB-LTD recordings, a pair of EPSCs (50 ms interval) were evoked at 0.05 Hz and three successive EPSCs were averaged and quantified relative to the normalized baseline. For DHPG mediated eCB-LTD experiments, cells were slightly depolarized to -50 mV, and DHPG (50 µM) was added to the perfusion following a baseline period^32,45^. For WIN mediated LTD experiments, WIN (2 µM) was added to the perfusion. In various control experiments, AM251 (10 µM) was added to the perfusion to block CB1Rs. Paired-pulse ratios were measured by dividing the peak amplitude of the second evoked EPSC by the first EPSC.

### DSI recordings

For DSI experiments, a high-chloride internal solution was used including: 125.2 mM CsCl, 10 mM NaCl, 10 mM HEPES, 1 mM EGTA, 2 mM QX-314 chloride, 0.1 mM CaCl_2_, 4 mM Mg-ATP, 0.3 mM Na_3_-GTP, and 8 mM disodium phosphocreatine (280-290 mOsm, pH 7.3 with CsOH). NBQX (10 µM) and R-CPP (10 µM) were included in the perfusion to block AMPAR- and NMDAR-mediated currents respectively. In DSI experiments measuring sIPSC charge, high-Ca2+ ACSF was used (4 mM Ca2+, 0.5 mM Mg2+) to increase the rate of spontaneous events, and sIPSCs were recorded for a baseline of 60 seconds before depolarization to 0 mV for 5 seconds and additional recording of sIPSCs for 60 seconds after depolarization^37^. sIPSC charge (integrated current) was binned every 2 seconds, normalized to the average of the 10 seconds (5 bins) preceding depolarization, and the normalized charge before depolarization, after depolarization, and 20 seconds after depolarization were compared. For DSI experiments measuring evoked IPSCs, a pair of evoked IPSCs (50 ms interval) were evoked at 0.2 Hz and average peak amplitude and average PPR were measured before, after, and 20 seconds after depolarization. 3 traces were averaged / cell.

### AEA-loading LTD recordings

For AEA-loading experiments, AEA (50 µM) was included in the internal solution as previously described^52,53^. Briefly, evoked EPSCs were recorded starting 5 minutes after achieving whole-cell configuration in order to allow EPSC amplitudes to stabilize. Baseline periods were measured in the 5-10 minute period following whole-cell break in, and all peak amplitudes were normalized to the average EPSC amplitude during this baseline period.

### Viral plasmid constructions

To generate pAAV-hSyn-GFP-IRES-mSNCA/hSNCA plasmids, GFP, IRES and SNCA coding sequences were cloned and sequentially stitched using overlapping PCR. Then, GFP-IRES-mSNCA/hSNCA fragments were digested with *Age*I/*Nhe*I and inserted into a pAAV-hSyn-Empty plasmid (Ding lab collection). Specifically, hSNCA and mSNCA were amplificated using pTB-hSyn-hSNCA and pTB-hSyn-mSNCA plasmids (gifts from Sudhof lab) as templates, respectively. To truncate mSNCA, a pair of primers were used to amplificate the coding sequence of 1-95 amino acids of the mSNCA. Then, full-length mSNCA was removed by XbaI/NheI digestion and replaced by mSNCA (1-95) to generate the pAAV-hSyn-GFP-IRES-mSNCA(1-95) plasmid. Similarly, we introduced S129A, S129D, A11P/V70P or A30P mutations into pAAV-hSyn-GFP-IRES-hSNCA construct by replacing the wild type hSNCA with corresponding mutants. All mutants were subcloned from pCMV5-hSNCA mutant plasmids (gifts from Sudhof lab). For viral packaging, all plasmids were prepared using EndoFree Plasmid Maxi Kit (Qiagen, Cat No.12362). For the TeNT lentivirus, a FUW-UBC-EGFP-2A-TeNT plasmid (gift from Sudhof lab) was used.

### Viral packaging

All SNCA viruses were packaged into AAV8 capsid and purified by discontinuous iodixanol gradients and ultracentrifugation as previously described^64^. Briefly, 640 ul (1mg/ml) Polyethylenimine Hydrochloride (PEI) solution (MW 40 kDa, pH7.0, Cat 24765) was mixed with serum-free DMEM media containing 3 ug of AAV genome plasmid, 35 ug of AAV8 capsid plasmid (AAV8-Rep/Cap) and 100 ug of helper plasmid (pHGTI-adeno1), and incubated at RT for 15 min. Then, DNA/PEI mixture was slowly added into 293T cell culture (5x 15 cm dishes) and mixed well. After incubation with 293T cells at 37 °C for 24 h, transfection media was replaced with fresh serum-free DMEM. 72 h after transfection, culture media was harvested and filtered through 0.44 um filters to get rid of cells and debris. To precipitated virus, collected media was incubated with 0.4 M NaCl and 8.5% PEG8000 at 4 °C for 1.5 hours followed by spinning down at 7000 g for 10 min. Viral particles were resuspended with 10ml lysis buffer (150 mM NaCl, 20 mM Tris, 10 mM MgCl_2_, pH 8.0), then incubated with 25 U/ml Benzonase (Sigma, E8263) at 37°C for 10 min. Crude virus isolate was then transferred to the top layer of a iodixanol step gradient (15%, 25%, 40%, and 60%) and centrifugated at 46,500 rpm (Beckman VTi50 rotor) for 90 minutes at 4°C. Purified viruses were collected from a new formed layer between 40% and 60% layers after centrifugation, washed twice with PBS and concentrated using Amicon Ultra-15 centrifugal filter units (100 kDa, EMD Millipore Cat#UFC10008). Viruses were aliquoted and stored in -80 °C. 5ul of virus was resolved by SDS-PAGE gel for purity assessment and semi-quantitative titration. TeNT lentivirus was prepared by the Stanford Gene Vector and Virus Core.

### Stereotaxic viral injections

Stereotaxic injections of AAVs and lentiviruses were performed on male and female adult mice (3-months old) under isoflurane anesthesia. A total volume of 100-300 nL was injected unilaterally into the left dorsal striatum (from bregma, AP: 1.0, ML: 2.4, DV: 3.4). Injections were performed using a micropipette (VWR) pulled with a long, narrow tip size (∼10-20 µm) using a micropipette puller (Sutter Instruments). Glass micropipettes were slowly inserted into the brain and left for 10 minutes before virus was injected at an infusion rate of 100 nL / min. The pipettes were then slowly retracted 10 minutes after infusion, and animals were sutured and monitored post-surgery. Acute brain slice recordings were performed 2-6 weeks following injections, where infected cells were identified by GFP fluorescence (BX51, Olympus).

### 2-photon imaging of GRAB_eCB2.0_

After 4 weeks following stereotaxic injection (see above) of AAV9-GRAB_eCB2.054_, acute brain slices were prepared (see above) for imaging. Two-photon imaging was performed using a custom-modified Olympus microscope (FV1200) with a Mai Tai Ti:sapphire laser (Spectra-Physics) with a low laser power (output optical power <40 mW) to avoid phototoxicity, and a 25x/1.05 NA water-immersion objective. A 920-nm wavelength was used to excite the GRAB_eCB2.0_ sensor, and fluorescence was collected using a 495–540-nm filter. Electrical stimulation consisted of 10 pulses (0.2 ms duration) delivered at 20 Hz (Dong et al., 2020). Pharmacological experiments included addition of 10 µM AEA and/or 10 µM AM251 to the ACSF perfusion at 2-3ml/min. All images were acquired at a frame rate of 2 Hz with a resolution of 512×512 pixels. The average pixel intensity of each frame was quantified and normalized to the baseline intensity (average intensity of first 4 frames [2 seconds] before stimulation) to quantify GRAB_eCB2.0_ sensor activity and response to pharmacology.

### Immunocytochemistry

In a subset of recordings, the brain slices were fixed by transferring to wells of 4% paraformaldehyde in 0.1 M Phosphate Buffer (0.1 M PB, pH 7.4) overnight at 4°C. Slices were then washed in PBS 3 times (10 minutes each) at room temperature (RT), before being mounted using an antifade mounting medium including a nuclear DAPI stain (VECTOR, USA). For α-Syn staining experiments (e.g. Fig. 3), fixed slices were washed in PBS 3 times (10 minutes each) before undergoing a block incubation with 2% bovine serum albumin and 10% normal donkey serum in PBS with 0.5% Triton X-100 (Sigma; PBS-T) (1 hr, RT) to reduce non-specific binding. Slices were then incubated in a primary antibody solution containing antibodies against α-Syn (1:1000 dilution, BD #610786) and GFP (1:100, ab5450) overnight at 4°C, followed by secondary antibodies conjugated to Alexa 647 (α-Syn), Alexa 488 (GFP), and Alexa 555 (biocytin-filled cells) (1 hr, RT) before washing and mounting. Images were acquired using a confocal microscope (Leica DM2500) with consistent settings used across all slices.

## QUANTIFICATIONS AND STATISTICAL ANALYSIS

### Statistics

Repeated measurements (e.g., input-output curves, repeated-stimulation release dynamics, etc.) were analyzed using 2-way repeated measures ANOVA with post-hoc tests. All two-sample comparisons (e.g., LTD comparisons, PPRs, etc.) were analyzed with nonparametric tests (Mann-Whitney or Wilcoxon tests). Unless otherwise specified, data is presented as mean ± SEM (standard error of the mean), with all statistical tests, statistical significance values, and sample sizes described in the figure legends. Statistical thresholds used: * p < 0.05, ** p < 0.01, *** p < 0.001, **** p < 0.0001, ns: non-significant.

## DATA AND CODE AVAILABILITY

Electrophysiology and imaging datasets have not been deposited in a public repository but are available from the corresponding authors upon request.

---

**Extended Data Figure 1.**
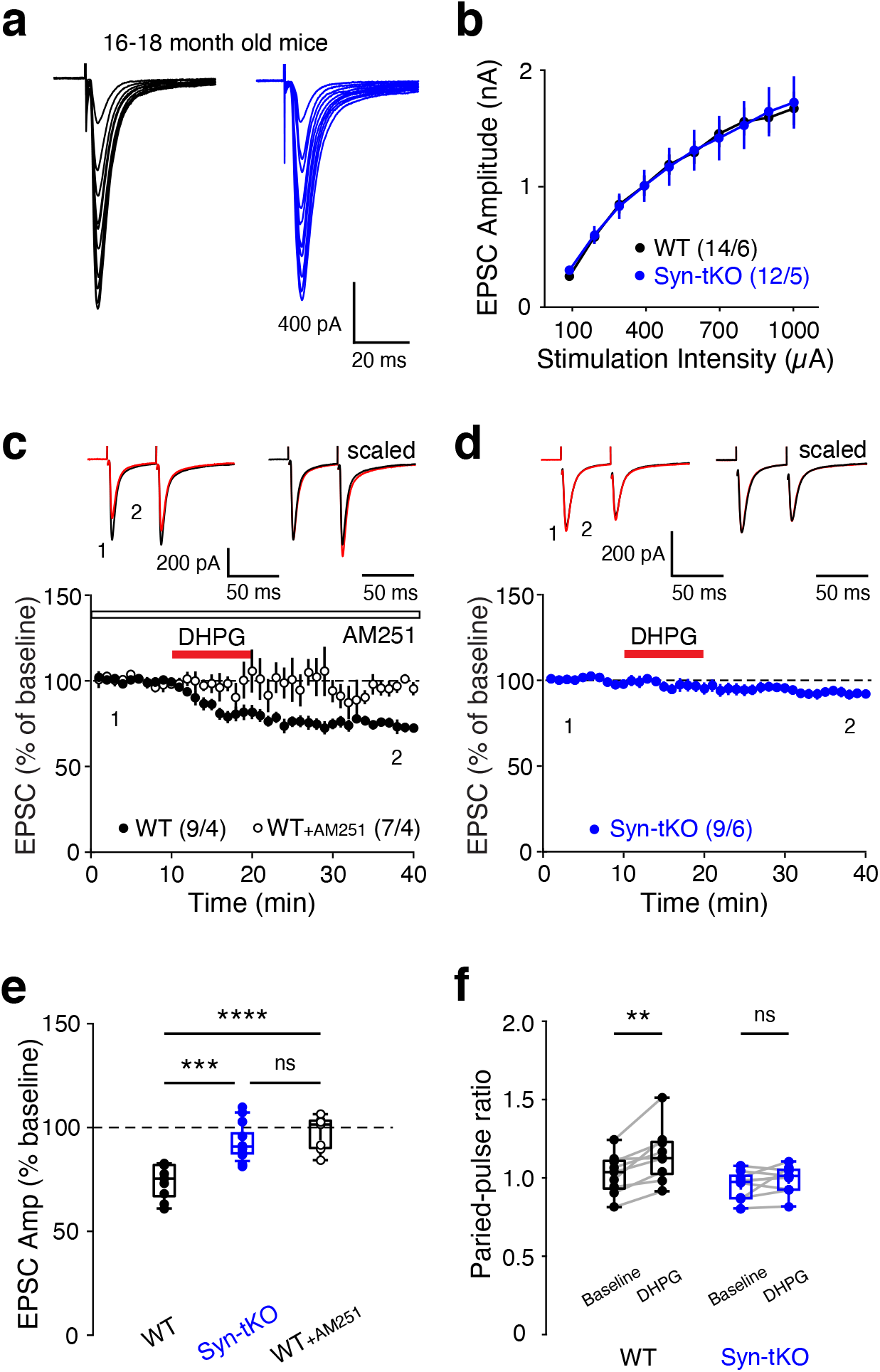
Normal corticostriatal input-output curves but impaired eCB-LTD in aged Syn-tKO mice. (**a)**, Representative traces of evoked corticostriatal EPSCs in WT and Syn-tKO SPNs from aged mice (16-18 months old) across a range of stimulation intensities. (**b)**, Normal input-output curves in Syn-tKO mice (WT: n = 14 cells / 6 mice; Syn-tKO: n = 12 cells / 5 mice; p = 0.960). (**c-e**), Summary of DHPG-mediated eCB-LTD in aged (16-18 months old) WT mice (**c**), which is fully blocked by the CB1R antagonist AM251 (10 µM) (WT: n = 9 cells / 4 mice, 73.88 ± 2.74%; WT + AM251: n = 7 cells / 4 mice, 97.06 ± 3.20%; p = 3.026e-5); top, representative traces. (**d**) eCB-LTD is impaired in aged Syn-tKO mice compared to WT (Syn-tKO: n = 9 cells / 6 mice, 92.57 ± 2.57%; p = 1.955e-4); top, representative trace. (**f**), Significant increase in aged WT PPRs (baseline: 1.02 ± 0.04; post-DHPG: 1.15 ± 0.06; p = 3.9e-3) but not in aged Syn-tKO PPRs (baseline: 0.95 ± 0.03; post-DHPG: 0.99 ± 0.03; p = 0.301). Data are mean ± s.e.m. Statistical significance was assessed by 2-way repeated measures ANOVA (**b**), ANOVA with multiple comparisons (**e**), and Wilcoxon signed tests (**f**) (**** p < 0.0001; *** p < 0.001; ** p < 0.01; n.s. non-significant).

**Extended Data Figure 2.**
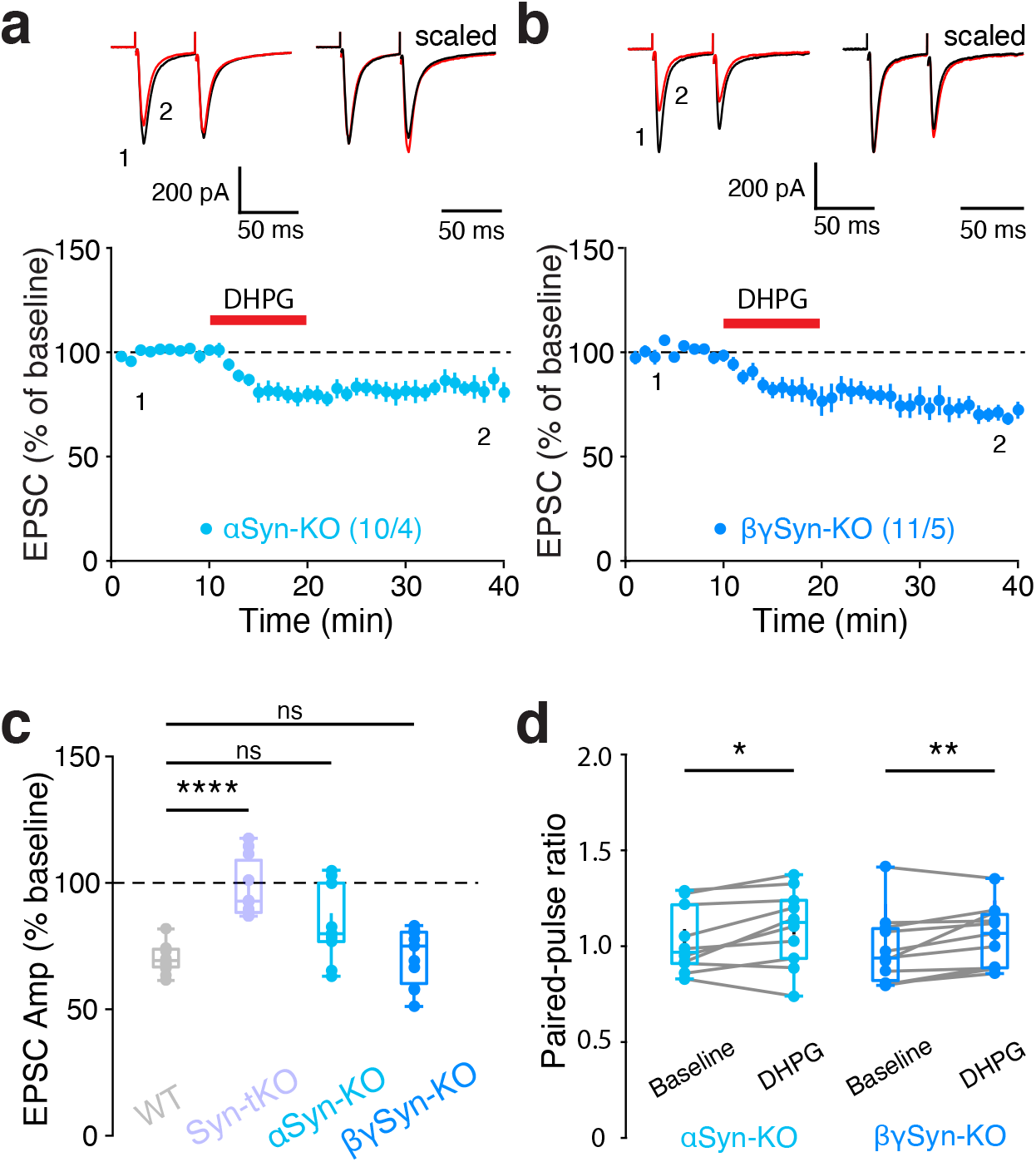
Normal DHPG-LTD in α-Syn-KO and βγ-Syn-KO mice. (**a-c**), DHPG-mediated eCB-LTD is normal in (**a**) α-Syn-KO (WT data from Fig. 1: n = 11 cells / 4 mice, 69.95 ± 1.70%; α-Syn-KO: n = 10 cells / 4 mice, 83.44 ± 4.67%; p = 0.162) and (**b**) βγ-Syn-KO mice (n = 11 cells / 5 mice, 71.00 ± 3.38%; p = 0.957). (**d**), Significant increases in PPR in both α-Syn-KO (baseline: 1.03 ± 0.05; post-DHPG: 1.10 ± 0.06; p = 0.037) and βγ-Syn-KO mice (baseline: 0.98 ± 0.06; post-DHPG: 1.05 ± 0.05; p = 9.8e-3). Data are mean ± s.e.m. Statistical significance was assessed by ANOVA with multiple comparisons (**c**), and Wilcoxon signed tests (**d**) (**** p < 0.0001; ** p < 0.01; * p < 0.05; n.s. non-significant).

**Extended Data Figure 3.**
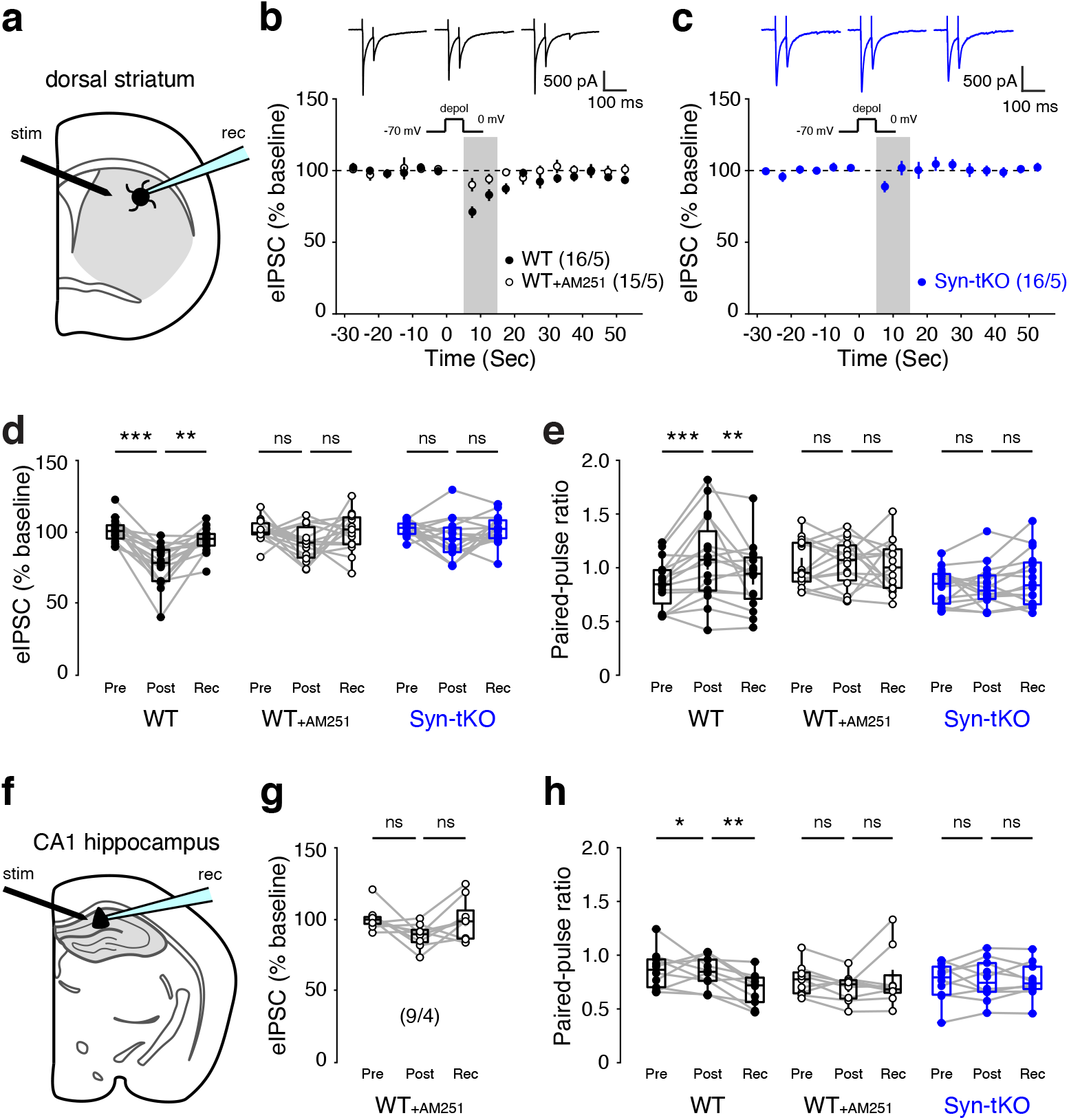
DSI is impaired in the dorsal striatum and CA1 of hippocampus of Syn-tKO mice. (**a**), Schematic of evoked DSI experiments in the dorsal striatum. (**b-d**), Summary of DSI (evoked IPSCs) in recorded SPNs in the dorsal striatum of WT mice (n = 16 cells / 5 mice; pre-depol: 100.94 ± 2.09%; post-depol: 76.95 ± 3.64%; recovery: 94.43 ± 2.16%; p = 4.378e-4, p = 7.764e-4), which is blocked by AM251 (10 µM) (n = 15 cells / 5 mice; pre-depol: 101.15 ± 1.99%; post-depol: 92.13 ± 3.14%; recovery: 99.97 ± 3.50%; p = 0.055, p = 0.277). Striatal DSI is impaired in Syn-tKO mice (**c, d**) (n = 16 cells / 5 mice; pre-depol: 102.19 ± 1.29%; post-depol: 95.25 ± 3.54%; recovery: 101.64 ± 2.53%; p = 0.134, p = 0.134). (**e**), Significant transient increases in PPR during striatal DSI in WT cells (pre-depol: 0.85 ± 0.06; post-depol: 1.08 ± 0.10; recovery: 0.92 ± 0.07; p = 2.3e-3, p = 0.030) but not in WT cells in the presence of AM251 (pre-depol: 1.04 ± 0.06; post-depol: 1.04 ± 0.06; recovery: 1.01 ± 0.06; p = 0.847, p = 0.600) or in Syn-tKO cells (pre-depol: 0.83 ± 0.04; post-depol: 0.82 ± 0.05; recovery: 0.89 ± 0.06; p = 0.959, p = 0.134). (**f**), Schematic of evoked DSI experiments in CA1 of the hippocampus. (**g**), Summary of DSI in recorded principal neurons in CA1 of WT mice in the presence of AM251 (10 µM) (n = 9 cells / 4 mice; pre-depol: 101.17 ± 2.82%; post-depol: 89.22 ± 2.67%; recovery: 100.26 ± 4.74%; p = 0.055, p = 0.098). (**h**), Significant transient increases in PPR during hippocampal DSI in WT cells (pre-depol: 0.69 ± 0.06; post-depol: 0.84 ± 0.05; recovery: 0.69 ± 0.05; p = 0.037, p = 5.9e-3) but not in WT cells in the presence of AM251 (pre-depol: 0.77 ± 0.05; post-depol: 0.70 ± 0.04; recovery: 0.78 ± 0.09; p = 0.098, p = 0.570) or in Syn-tKO cells (pre-depol: 0.75 ± 0.06; post-depol: 0.77 ± 0.06; recovery: 0.77 ± 0.05; p = 0.770, p = 1.000). Data are mean ± s.e.m. Statistical significance was assessed by Wilcoxon signed tests (**d, e, g, h**) (*** p < 0.001; ** p < 0.01; * p < 0.05; n.s. non-significant).

**Extended Data Figure 4.**
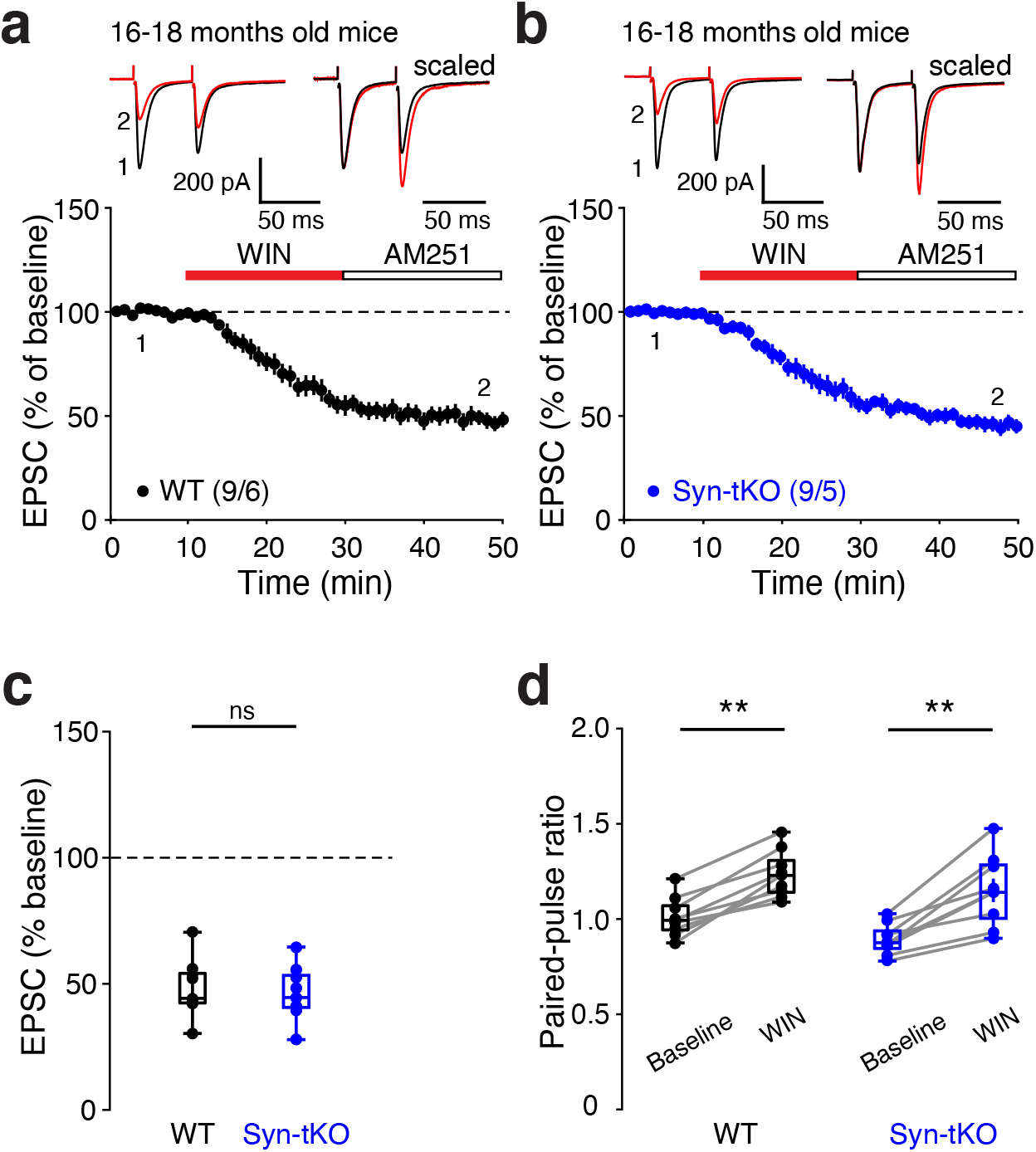
Normal WIN-LTD in aged Syn-tKO mice. (**a-c**), WIN application results in indistinguishable corticostriatal LTD in aged (16-18 months) WT and aged Syn-tKO mice (WT: n = 9 cells / 6 mice, 48.11 ± 3.78%; Syn-tKO: n = 9 cells / 5 mice, 46.01 ± 3.56%; p = 0.605). (**d**), Significant increases in PPRs in both aged WT (baseline: 1.02 ± 0.03; post-WIN: 1.24 ± 0.04; p = 3.9e-3) and aged Syn-tKO mice (baseline: 0.90 ± 0.03; post-WIN: 1.16 ± 0.06; p = 3.9e-3). Data are mean ± s.e.m. Statistical significance was assessed by Mann-Whitney (**c**) and Wilcoxon signed tests (**d**) (** p < 0.01; n.s. non-significant).

**Extended Data Figure 5.**
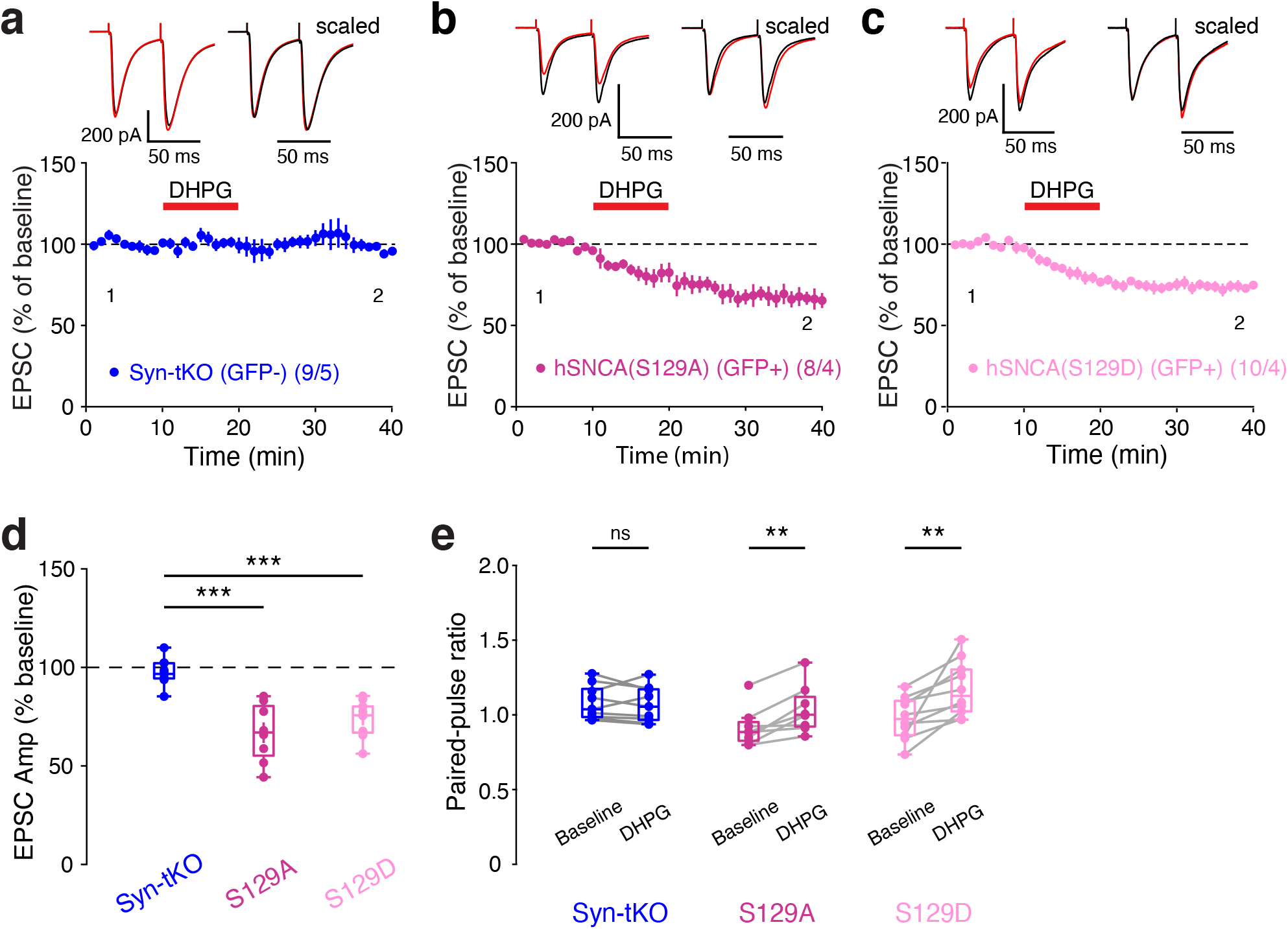
α-Syn S129A (phospho-deficient) and α-Syn S129D (phos-phor-mimic) mutations do not disrupt viral α-Syn rescue of eCB-LTD. (**a-d**), Neither S129A (b) nor S129D (**c**) mutations affected the ability of postsynaptic viral α-Syn to rescue eCB-LTD (GFP-, pooled: n = 9 cells / 5 mice, 97.54 ± 2.28%; GFP+, S129A: n = 8 cells / 4 mice, 66.89 ± 5.24%; p = 1.03e-5; GFP+, S129D: n = 10 cells / 4 mice, 73.62 ± 2.89%, p = 1.39e-4). (**e**), Significant PPRs observed in cells infected with S129A α-Syn (baseline: 0.91 ± 0.05; post-DHPG: 1.04 ± 0.06; p = 7.8e-3) and S129D α-Syn (baseline: 0.98 ± 0.04; post-DHPG: 1.18 ± 0.06; p = 2.0e-3), but not in uninfected cells (baseline: 1.08 ± 0.04; post-DHPG: 1.07 ± 0.04; p = 0.496). Data are mean ± s.e.m. Statistical significance was assessed by ANOVA with multiple comparisons (**d**) and Wilcoxon signed tests (**e**) (*** p < 0.001; ** p < 0.01; n.s. non-significant).

**Extended Data Figure 6.**
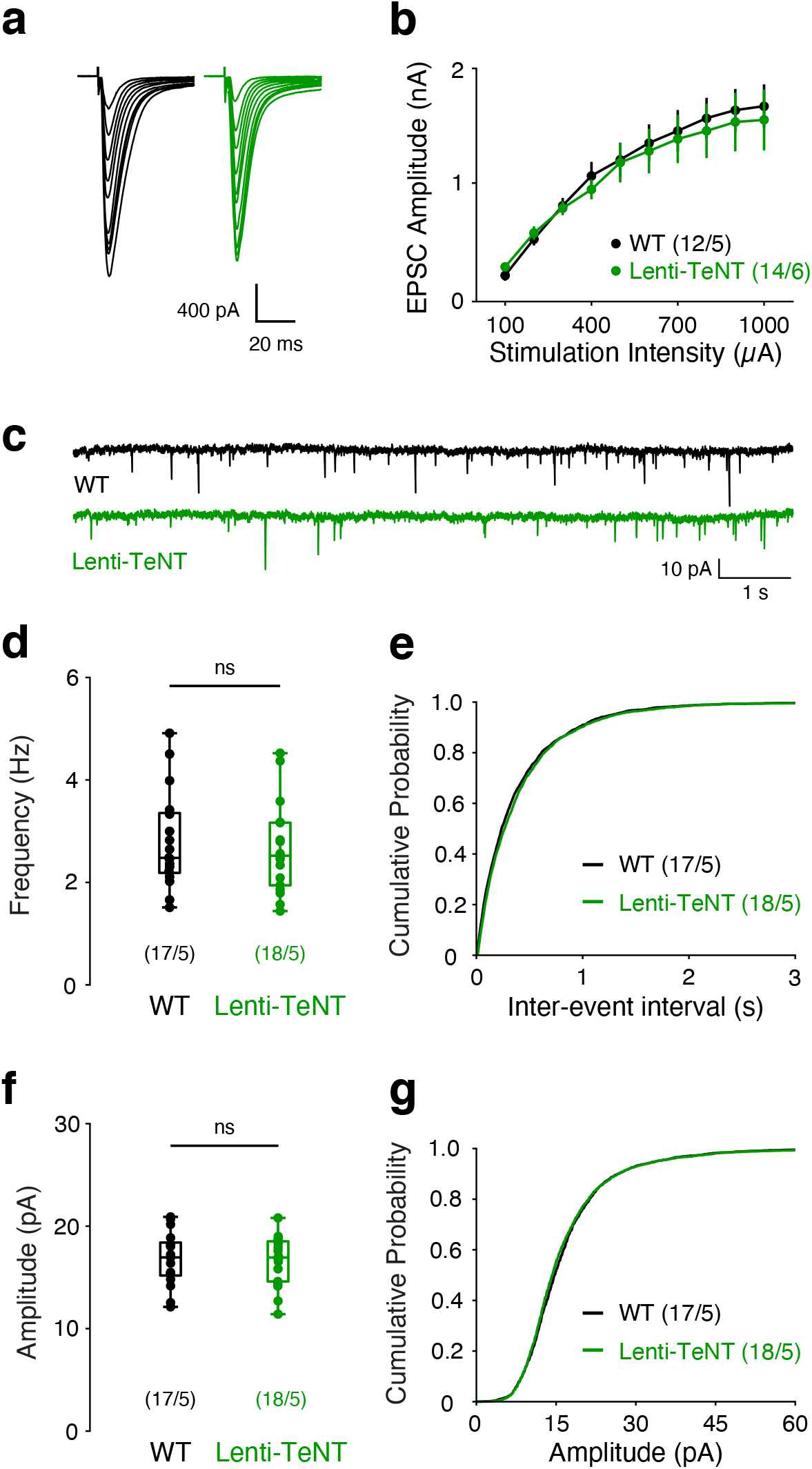
Lentiviral TeNT does not disrupt basal synaptic properties of striatal SPNs. (**a**), Representative traces of evoked corticostriatal EPSCs in WT (GFP-) and TeNT-infected (GFP+) SPNs across a range of stimulation intensities. (**b**), Normal input-output curves in TeNT-expressing SPNs (GFP-: n = 12 cells / 5 mice; GFP+: n = 14 cells / 6 mice; p = 0.787). (**c**), Representative traces of mEPSC recordings from WT (GFP-) and TeNT-expressing (GFP+) cells. (**d-g**), Lentiviral TeNT does not result in a change in (**d, e**) mEPSC frequency (GFP-: n = 17 cells / 5 mice, 2.81 ± 0.22 Hz; GFP+: n = 18 cells / 5 mice, 2.64 ± 0.20 Hz; p = 0.680) or in (**f, g**) mEPSC amplitude (GFP-: 16.78 ± 0.62 pA; GFP+: 16.65 ± 0.55 pA; p = 0.987). Data re mean ± s.e.m. Statistical significance was assessed by 2-way repeated measures ANOVA (**b**) and Mann-Whitney tests (**d, f**) (n.s. non-significant).

